# Neural activity flows through cortical subnetworks during speech production

**DOI:** 10.1101/2025.06.20.660783

**Authors:** Gregg A. Castellucci, Mac MacKay, Christopher K. Kovach, Farhad Tabasi, Jeremy D.W. Greenlee, Michael A. Long

## Abstract

Speech production entails several processing steps that encode linguistic and articulatory structure, but whether these computations correspond to spatiotemporally discrete patterns of neural activity is unclear. To address this issue, we used electrocorticography to directly measure the brains of neurosurgical participants performing an interactive speech paradigm. We observed a broad range of cortical modulation profiles, and subsequent clustering analyses established that responses comprised distinct classes associated with sensory perception, planning, motor execution, and task-related suppression. These activity classes were also localized to separate neural substrates, indicating their status as specialized networks. We then parsed dynamics in the planning and motor networks using unsupervised dimensionality reduction, which revealed subnetworks that were sequentially active throughout preparation and articulation. These results therefore support and extend a localizationist model of speech production where cortical activity ‘flows’ within and across discrete pathways during language use.

## INTRODUCTION

During speech production, an array of planning and motor processes is hypothesized to enable the translation of abstract thought to vocal tract action (e.g., conceptualization, lexical access, programming of articulatory movements)^1–3^. These operations have been hypothesized to represent separate preparatory and execution stages^4–6^ that possess divergent neuroanatomical substrates^7–16^. However, this notion remains controversial, as some behavioral and functional studies^17–23^ suggest that continuous and widespread neural processing underlies speech production. Such an interpretation is consistent with broader trends in systems neuroscience emphasizing the potential importance of distributed rather than localized dynamics^24–27^. Consequently, the neural architecture underlying this core human ability and the degree to which it consists of anatomically distinct processing units remains unclear.

Because speech planning and articulation unfold at subsecond timescales and are thought to involve many brain regions^6,15,21^, delineating the neural dynamics related to these behaviors necessitates precise temporal resolution as well as broad spatial coverage. These requirements are uniquely met by intracranial recordings in neurosurgical patients^28–31^, methods that we recently used to investigate the neural basis of interactive language use^32^. In this study, we found that different frontotemporal regions were preferentially active during periods of speaking in comparison to periods of planning, indicating that these general functions are indeed localized to different substrates. However, our statistical approach considered only a subset of responses and was not capable of testing whether the dynamics observed during speech production could be further subdivided on a functional or anatomical basis. Consequently, it is unknown whether these networks exhibit additional structure indicative of planning and motor subprocesses that are dissociable at a neural level.

Here we directly address this question by leveraging a large data set of intracranial recordings that offers extensive coverage of the left, right, and ventral cortical surfaces, which allows us to consider the full diversity of production-related responses observed during an interactive speech task. Using a combination of unsupervised analytical methods, we map the extent of neural responsivity, show that these dynamics form distinct brain networks, and track activity as it proceeds through a spatiotemporal sequence during planning and articulation. This investigation therefore provides strong evidence for a localized model of spoken language production and sheds light onto how large-scale cortical dynamics are coordinated to enable this complex behavior.

## RESULTS

### Neural activity during speech production forms functionally distinct clusters

Our first goal was to identify the range of cortical loci that were engaged during speech production. We used electrocorticography (ECoG) and stereoelectroencephalography to record neural activity from the cortical surfaces of 22 neurosurgical patients (n = 19 with left hemisphere coverage; Figure 1A, Table S1) as they performed a question-answer task featuring cognitive and sensorimotor demands similar to natural conversation^33–35^. Participants answered a battery of 20-119 critical information (CI) questions^32,33^ (mean ± standard deviation: 47 ± 23) that were read by an experimenter. In this paradigm, each question contains a critical word (i.e., the CI) that is necessary to plan a response; consequently, this task isolates the sensory phase of spoken interaction from periods of planning or motor function (Figure 1B) and permits neural dynamics to be associated with these three generalized processes^32,36^.

**Figure 1.**
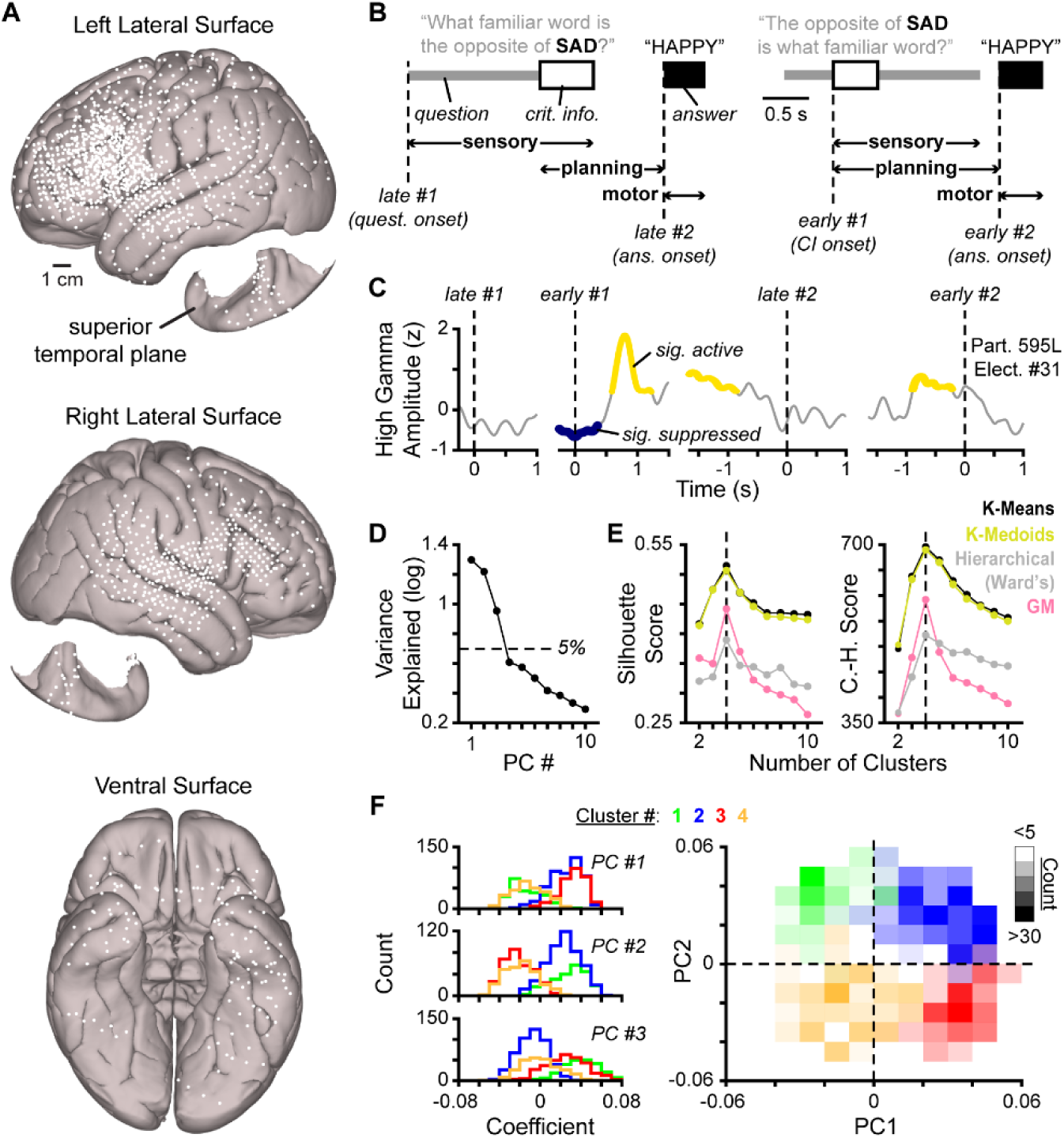
Four clusters of neural activity in an interactive language task. (A) Location of all analyzed electrodes shown on the cortical surface of a canonical brain. (B) Example trials of the critical information task where the critical word is presented either at the end (‘late’) or in the middle (‘early’) of the question; sensory, planning, and motor phases of each trial are indicated at bottom and task-related alignment points are denoted with dashed lines. (C) Average responses from an example electrode at each task-related alignment point with periods of significant activation and suppression indicated. (D) Amount of variance explained by each principal component (PC). (E) Empirical silhouette (left) and Calinski-Harabasz (C.-H.) (right) scores when clustering significantly modulated electrode responses in 3-dimensional PC space using four different clustering methods (indicated by line color); for k-means, k-medoids, and Gaussian mixture (GM) clustering, the median score across 500 iterations is reported. Dashed line indicates optimal clustering solution. (F) Distribution of PC coefficients for each response cluster depicted with histograms (left) and a heatmap (right); for the heatmap, color denotes cluster identity and transparency indicates the number of electrodes within each bin.

To assess brain function during task performance, we extracted the high gamma band of the intracranial recordings (70-150 Hz), as the amplitude of this signal is correlated with multiunit spiking and thus represents a readout of local neural activity^28–30^. We then identified recording sites whose high gamma activity was modulated at any of four nonoverlapping timepoints using a resampling procedure that controlled for behavioral differences between participants (Figure 1C, Figure S1). This method was developed to detect a broad variety of responses, and it revealed that 1,296 of 1,784 total electrodes (72.6%) were significantly activated and/or suppressed (see Methods) while answering CI questions. In comparison, our past work using this paradigm employed a GLM-based technique designed to identify sites that were significantly active throughout the duration of any task phase (i.e., Figure 1B), which indicated that 32.0% of electrodes were modulated in this fashion^32^. Therefore, our current statistical approach uncovers a diverse array of neural responses during task-based spoken interaction, and these dynamics are densely represented across the cortical surface.

We next tested if this widespread speech-related activity was organized into distinct categories or instead represented a continuum of dynamics. To do so, we assessed the similarity across the responses of all modulated electrodes using principal component analysis (see Methods). We found that only the first three principal components explained more than 5% of the variance (Figure 1D; Figure S2A) and that neural responses in 3-dimensional PC space were optimally organized into four clusters when assessed by multiple algorithms (Figure 1E,F); crucially, this clustering was not present when responses were shuffled (Figure S2B-C). Therefore, although we find that large-scale cortical activity supports CI task performance, these dynamics do not represent a continuum of sensorimotor processing^24^ – which would be denoted by a single distribution rather than functionally distinct groups.

### Neural response classes track task-relevant behaviors

We next aimed to determine whether the identified neural activity clusters reflect specific aspects of task engagement. To do so, we split electrodes into classes according to the optimal k-means clustering solution and examined group-level responses when aligned to behaviorally relevant timepoints. We found that individual electrodes exhibited considerable heterogeneity in their response profiles (Figure 2A); however, dynamics within a class corresponded to consistent patterns of modulation during the task (Figure 2B) while randomized electrode responses did not (Figure S2D,E). Specifically, Class #1 was active after question onset but rapidly suppressed after question offset (n = 236, 18.2% of modulated electrodes), Class #2 was active following CI onset and before the spoken answer (n = 473, 36.5%), Class #3 was active during and immediately prior to the answer (n = 315, 24.3%), and Class #4 was weakly modulated but generally suppressed between CI presentation and answer onset (n = 272, 21.0%) (Figure 2C). Furthermore, Classes #1-#3 were primarily active during the time windows where sensory, planning, and motor activity was expected based on task structure (Figure 2D), and each response class was largely active during a single phase of the CI task but inactive or suppressed in others (Figure S3A-C). In sum, these results indicate that speech-related sensory, planning, and motor function is represented in the brain by distinct neural dynamics (Classes #1-#3, respectively) and identify a pronounced type of task-related suppression during spoken interaction (Class #4).

**Figure 2.**
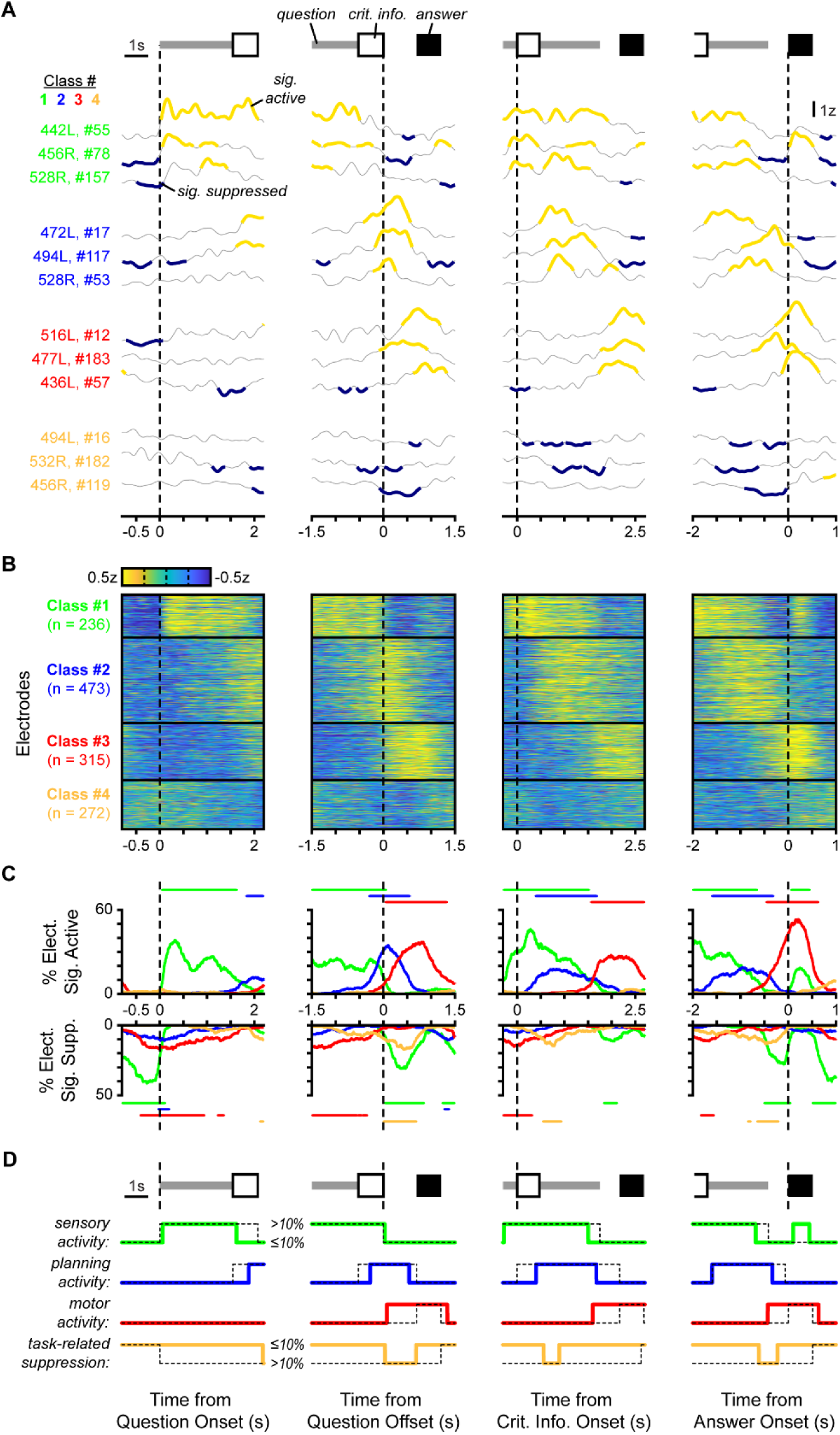
Neural response classes reflect the core phases of spoken interaction. (A) Average responses from example electrodes aligned to four different points in the critical information task; trial diagrams at top are normalized to the median trial duration across participants (see Methods). (B) Average responses for all electrodes in each activity class when aligned to the points in (A). (C) For the alignment points in (A), the percentage of electrodes from each class that display either significantly elevated (top) or suppressed (bottom) activity over time; thin bars at top and bottom indicate when more than 10% of electrodes within a class are significantly activated or suppressed, respectively. (D) For the same alignment points, the time windows where sensory activity, planning activity, motor activity, and task-related suppression is expected (dashed lines) and when more than 10% of electrodes within selected activity classes are significantly modulated (thick lines; smoothed with a 50-ms moving median window).

### Neural response classes are localized to distinct cortical loci

Our next goal was to test if response categories – as defined by our new unsupervised approach – were organized into spatially distinct networks or distributed broadly across the brain. We first examined individual participants and observed clear neuroanatomical clustering according to response class (Figure 3A), with electrodes typically neighboring (i.e., closer than 1 cm) electrodes from the same group (Figure 3B). To quantify this effect, we shuffled class identity across electrodes on a within-participant basis and found that the observed spatial structuring never arose randomly (Figure 3C); this approach likewise revealed that response classes were localized to consistent cortical areas across individuals when all recording sites were coregistered to a reference brain (Figure 3D). Therefore, the functional distinction between response classes is also reflected by anatomical segregation.

**Figure 3.**
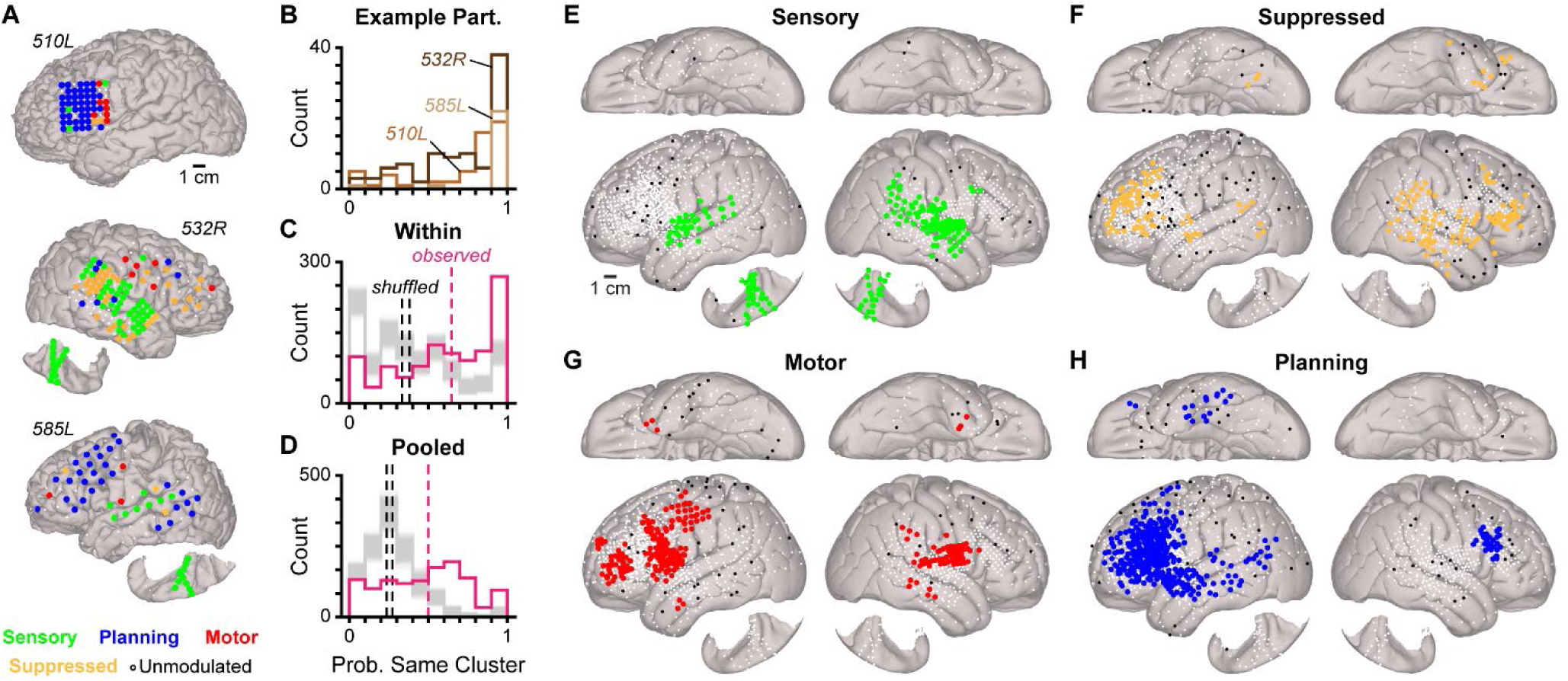
Neural response classes are localized to different anatomical substrates. (A) Location of all modulated electrodes in three example participants; electrode color indicates response class. (B) For each electrode within the participants presented in (A), the proportion of neighboring electrodes (i.e., located within 1 cm) from the same response class. (C-D) The same measure in B, but for all modulated electrodes when considering electrodes only from the same participant (C) or all electrodes pooled across participants (D); pink and black dashed lines denote the median value for the observed distribution and the range of median values when class labels are shuffled 1,000 times, respectively, with thin grey lines depicting the distribution for each iteration of shuffling. (E-H) Location of all sensory (E), suppressed (F), motor (G), and planning (H) class electrodes on canonical cortical surfaces; large colored points denote electrodes within voxels with significantly more electrodes of a given class than expected by chance (p < 0.05, permutation test) while small points denote significantly modulated electrodes outside these voxels and white points denote unmodulated electrodes or electrodes in other classes.

These results are important because they demonstrate that the brain networks underlying speech planning and motor function are dissociable from those associated with perception and task-related suppression – even when the diverse range of activity profiles observed during interaction is considered. To delineate these networks, we used a spatial permutation test to detect areas of the reference cortical surface where responses of each class were significantly overrepresented (p < 0.05; see Methods) while controlling for differences in recording coverage across participants (e.g., Figure 3A). As an internal control, we first used this method to define the speech-related sensory network as it has been extensively described in previous intracranial research^32,37–40^. We found that this network was largely restricted to bilateral temporo-parietal areas implicated in audition and speech perception^37–48^, including superior temporal gyri (STG) and transverse temporal gyri, but was more extensive in the right hemisphere (Figure 3E; Figure S3D). These findings demonstrate that sensory dynamics during spoken interaction are localized to established structures and indicate that our approach can effectively map language-associated cortical regions.

We next defined the suppressed network; while the relevance of this activity class during the CI task is unclear, its substrates have not been described previously. We found that this network extended widely across the right and left hemispheres but was most densely represented in right frontal and temporoparietal cortex as well as a subregion of left rostral middle frontal gyrus (MFG) (Figure 3F; Figure S3E). This class of cortical dynamics is therefore widely observed across the right lateral cortical surface but is also present in a more limited region of the left hemisphere.

Finally, we examined the networks underlying spoken language production. We found that the motor network was primarily situated within bilateral precentral gyrus (PrCG) and postcentral gyrus, which contain speech-related ventral and middle sensorimotor cortices^49–55^, as well as a left-lateralized prefrontal locus located within pars triangularis and rostral MFG (Figure 3G; Figure S3F). In contrast to sensorimotor and suppressed responses, the planning network was heavily left lateralized and encompassed frontotemporal cortical areas (Figure 3H; Figure S3G) that were previously implicated in planning using the CI task^32,36^ – such as IFG^9,10,56^, MFG^57–59^, anterior PrCG^5,60^, and anterior STG^6,10,61,62^ – in addition to planning-linked structures within left superior frontal^5,11,63,64^, fusiform^65,66^, and middle temporal gyri^6,41^. This network also included a subregion of right pars opercularis and caudal MFG, indicating that a restricted area of frontal cortex in the non-dominant hemisphere may be involved in language-related planning. In summary, our large intracranial data set enables us to perform a comprehensive mapping of the human speech production system.

### Neural activity flows across planning and motor regions

Our results demonstrate that separate cortical networks underlie speech planning and motor function, indicating that preparation and execution are discrete in the brain. However, further processing stages are thought to exist^2–5,32,67^, and these functions can be grouped into the following categories: domain-general conceptualization, planning operations that program semantic and syntactic structure, lower-level encoding of phonological and phonetic features, and motor processes that control articulation. We therefore hypothesized that the heterogeneity observed across individual electrodes (e.g., Figure 2A,B) might correspond to further functional subdivision within the broader networks we identified. As an initial test of this prediction, we examined how neural activity progressed before and during speech in a participant implanted with a high-density ECoG grid that spanned frontotemporal language areas^5,6,32,61^ (Figure 4A,B); this revealed a spatiotemporal ‘flow’ of activity across regions (Figure 4C), which suggests the presence of production-related subnetworks with differing temporal characteristics.

**Figure 4.**
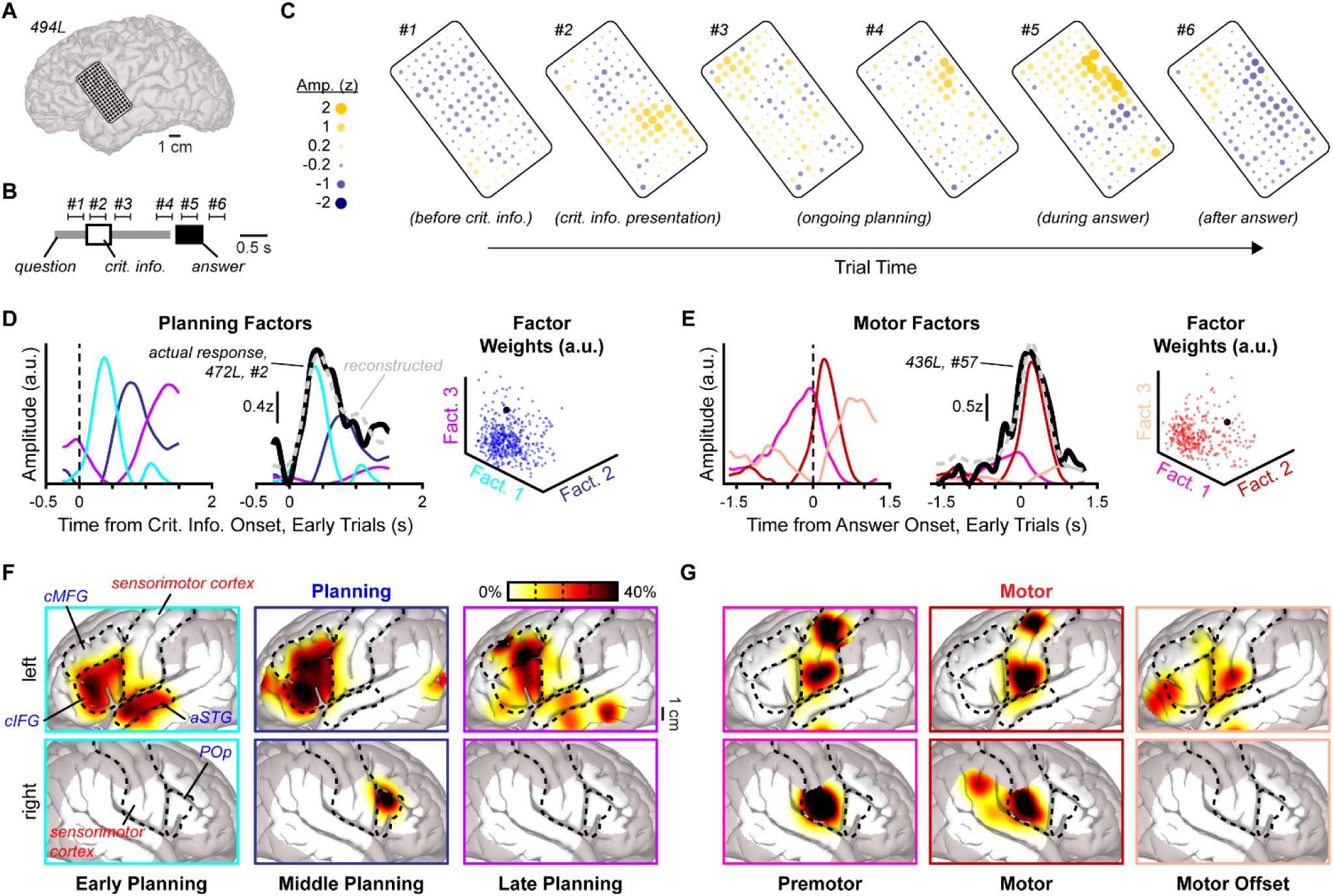
Nodes within the planning and motor networks exhibit distinct response profiles. (A) Location of high-density ECoG grid in an example participant. (B) Diagram of an early CI trial with analysis epochs numbered; trial is normalized to the median trial duration for the participant in (A) (see Methods). (C) Average activity within the 300 ms analysis windows numbered in (B) for all analyzed electrodes from the grid depicted in (A); activity in windows 1-3 and 4-6 were aligned to answer and critical information onset, respectively. (D-E) Component response profiles recovered with nonnegative matrix factorization in planning (D) and motor (E) class electrodes (see Methods and Figure S4) (left), with an example reconstructed electrode signal (middle), and the distribution of component weights (right); in the scatterplots, the example electrodes are depicted as black points. (F-G) The density of high-weight planning (F) and motor (G) electrodes within voxels that are weighted significantly higher for each component response profile than expected by chance (p < 0.05, permutation test); heatmaps are spatially smoothed with a 7.5 mm Gaussian kernel (see Methods) and brain areas labeled at top left in blue and red denote regions associated with planning and motor function, respectively.

To determine whether planning and motor dynamics could be further subdivided according to response timing, we used nonnegative matrix factorization – a dimensionality reduction technique that decomposes complex signals into component parts (i.e., factors)^68^ – to identify repeated temporal profiles embedded within the activity of individual electrodes (Figure S4A-C; see Methods). This analysis revealed that each response class was comprised of multiple sub-responses varying in timecourse and degree of representation across recording sites (Figure 4D,E; Figure S4D,E). In particular, planning-related modulation was composed of early, middle, and late components (Figure 4D, Figure S4F), while motor activity exhibited components that peaked either before speech onset, shortly after speech onset, or around the offset of the spoken answer (Figure 4E, Figure S4G). Different types of dynamics are therefore maximally engaged throughout planning and articulation, consistent with multiple preparatory and motor stages that are distinct at a neural level.

We next examined whether motor and planning response components were localized to specific areas within their respective networks. Because electrodes did not form discrete clusters according to their factor weights (Figure 4D,E, right; Figure S4D,E), we instead detected cortical regions that preferentially exhibited each response component using our spatial permutation approach (see Figure 3E-H). We found that many loci were weighted significantly higher for a component than would be expected by chance (see Methods) and individual components were often overweighted in different areas (Figure S4H,I). To visualize this effect, we computed the spatial density of high-weight electrodes (i.e., possessing a weight higher than the median value) within these regions; this uncovered a sequence of activity (1) propagated from left anterior STG and caudal IFG, to bilateral caudal IFG and MFG, and then to posterior aspects of left caudal IFG and MFG during planning (Figure 4F), (2) progressed to bilateral motor structures during premotor processing and articulation, and finally terminated in left ventral PrCG and prefrontal cortex (Figure 4G). These results show that individual planning and motor response components are preferentially represented at distinctive network ‘nodes’ – which may correspond to specialized processing centers for the cognitive and sensorimotor processes underlying speech.

## DISCUSSION

Using intracranial recordings, we examined how the neural systems underlying speech production are organized in the human brain. We found that widespread cortical modulation unfolds when speaking in an interactive context but that embedded within these dynamics are distinct networks associated with planning and articulation. These networks are functionally and anatomically separate from those linked to sensory processing and task-related suppression, indicating the existence of a dedicated pathway for spoken language generation. Furthermore, activity flows across different regions of the planning and motor networks before and during speech, supporting the idea that specific production-related operations (e.g., lexical access, articulatory programming) are performed within spatiotemporally discrete subnetworks.

### Support for a localizationist model of neural function

Because our data set offered wide and dense coverage of the cortical surface, we could explicitly test whether the range of neural responses observed during spoken interaction was organized according to localizationist or distributed models of brain function – which predict anatomically distinct categories of neural dynamics or a widely dispersed continuum of activity^25^, respectively. We found that responses were divided into distinct clusters with divergent spatial properties and thus determined that most of the brain appears to support the task performance only because activity is spread across multiple localized networks with specialized functions. These results are therefore inconsistent with recent hypotheses about global cortical processing during real-world language use in humans^23^ and behavioral engagement in model organisms^24,26,27^; instead, our results support localizationist accounts of language^5,6,41,69,70^ and modular frameworks of cognitive function^71–73^.

Neural processing during our interactive question-answer paradigm was specifically segregated into classes associated with perceiving the question (sensory), preparing a response (planning), or speaking an answer (motor). These response types were restricted to different areas of cortex, with sensory and motor dynamics localized to largely non-overlapping bilateral networks and planning activity largely restricted to left hemisphere frontotemporal areas – in agreement with classical views of lateralization for higher-level language function^70,74,75^. We also identified a fourth class of cortical dynamics that were primarily suppressed during planning, and this activity pattern was mainly observed in right cortical regions. The role of this response type is unknown, but similar suppression in the default mode network has been reported to occur during task performance^76^. We hypothesize that this phenomenon reflects the inhibition of regions unrelated to language and social behavior, which may minimize irrelevant inputs into the planning network.

### Importance of network organization for naturalistic speech production

The planning network identified in this study overlaps considerably with the previously described left hemisphere ‘language network’, which was defined using fMRI and non-interactive tasks^41,77^. However, a key property of this network is its apparent multimodality, as it displays elevated BOLD activity during both the generation and comprehension of speech^41^. In contrast, our planning network was largely inactive or even suppressed outside of planning periods^32^ (Figures 2 and S3), indicating that its principal role relates to production. These findings are supported by our previous perturbation experiments, which demonstrated that direct electrical stimulation of frontal planning areas during CI task performance regularly induces delayed responses and errors but rarely elicits sensory-related deficits^36^. Instead, disrupted speech perception has been shown to primarily result from stimulation of specialized temporal regions^39^ located within the sensory network (i.e., Figure 3E). The identified planning network therefore appears dedicated to preparatory aspects of speaking, at least in our interactive paradigm.

While the discrepancy between our findings and previous work on the language network may arise partly from technical differences (i.e., ECoG vs. fMRI), we hypothesize that the major contributor is the unique task demands imposed during interactive language use. Specifically, responding to CI questions – and a conversational partner more generally – with the subsecond latencies characteristic of spoken interaction^78,79^ requires a speaker to simultaneously plan their response while also perceiving their partner’s speech (i.e., Figure 1B, right)^34,80,81^. This behavior therefore requires a degree of multitasking that is largely absent from paradigms typically used to investigate the neural basis of language (e.g., naming, passive listening, repetition)^41,56,82,83^. Because perception and response preparation do not overlap in such tasks, we propose that the planning network can inherit epiphenomenal activity from the sensory network; in contrast, multitasking during our interactive experiments necessitates the planning network to operate concurrently and independently from the sensory network, which we suggest prevents the development of perception-related activity.

Natural face-to-face interaction in humans requires extensive multitasking, as interactants must monitor their partner’s speech, prosody, gestures, facial expressions, and gaze to appropriately plan and initiate responses consisting of the same multimodal behaviors^35,80,81,84–87^. Furthermore, the multiword speech produced during natural language use often requires simultaneous planning and articulation (i.e., incremental planning)^88–91^. We postulate that parallel processing modules are required to support these multitasking demands, and the substrates of these modules may be the distinct cortical networks we identified. Consistent with this hypothesis, our previous intracranial recordings during unconstrained conversation also revealed distinct clusters of neural activity^32^.

### Support for distinct planning stages during speech production

The planning and motor networks isolated in this study were comprised of several regions exhibiting a family of responses (e.g., Figure 2A,B). However, we found activity within a network is not homogenous – instead, dynamics could be decomposed into distinct sub-responses whose activity peaked at different times prior to and during speech, and these response components were preferentially localized to different regions. These results therefore provide important neural evidence for distinct production-related processing stages by demonstrating the existence of subnetworks possessing unique temporal characteristics. In agreement with this conclusion, previous studies have shown that individual regions of our planning network display activity correlated with different linguistic features during speech production^11,12,83^ and can evoke specific semantic, syntactic, and phonological deficits when perturbed^7–10,13,14,92^. Nevertheless, planning and motor response components were not organized categorically and could be observed to varying extents throughout their respective networks. This is consistent with recent single unit recordings in neurosurgical patients demonstrating that neuronal spiking within a cortical site is typically correlated with many linguistic properties^93,94^ and past work showing that activity in the fMRI-defined language network is associated with multiple syntactic and semantic variables^41^. In sum, our findings suggest that within-network activity possesses features of localized and distributed function, with information about each neural processing step shared across a network but most strongly encoded within specific cortical areas.

This study has important theoretical implications for understanding how planning unfolds while speaking. In particular, the existence of independent planning and motor networks indicates that high-level planning operations are separate from motor processing – a notion that is questioned by some speech production frameworks^2,17–19^. Furthermore, we found that activity within the motor network exhibits a premotor component, implying that late-stage planning operations (e.g., phonetic planning, articulatory programming) are performed by the motor system. This premotor activity was most strongly represented in PrCG, which has recently been hypothesized to participate in both speech motor planning and the execution of vocal tract movement^5,60,95^.

In summary, our results support a model where early planning processes (e.g., conceptualization, lexical access) occur within a network that is functionally distinct from the motor system, which then separately performs low-level premotor and motor operations. This hypothesis is supported by psycholinguistic experiments using the CI paradigm, which have found that semantic and phonological structure appears to be planned rapidly after CI onset^96^ but articulatory posturing is delayed until the end of the question^97^ – thus indicating that speech movements are not planned immediately after phonological encoding in this task.

### Conclusion and future opportunities

This study demonstrates that interactive speech production is enabled by multiple planning and motor processes with distinct spatiotemporal representations in the brain, thus providing key insights into the neural architecture that underlies spoken language. Further investigation is now needed to determine the specific mental processes performed within the production-related networks and subnetworks our work has uncovered. This topic is well-suited to investigation using direct electrical stimulation paired with more sophisticated behavioral analysis. For example, stimulating the subnetwork we found to be preferentially active during “early planning” might result in a particular type of high-level errors (e.g., semantic errors, syntactic errors) while disordering activity in other planning regions could lead to different effects (e.g., phonological errors, speech arrest). Additionally, because our methods require averaging across a set of highly controlled behavioral data, it is unclear how the identified networks are organized in individual brains. Further analysis of raw neural signals from single participants with high spatiotemporal resolution (e.g., Figure 4A) will be critical for establishing the number of networks/subnetworks that exist, defining the nature of anatomical separation between networks, and understanding how network structure generalizes across the population.

In conclusion, this study provides foundational information about how cortical networks are organized to support speech production – thus offering an important mesoscale complement to recent single-neuron analyses of language function within single recording sites^43,93,94^. By characterizing the functional and anatomical relation between planning and motor processes, we hope to inform future work on brain machine interfaces. These efforts have made groundbreaking progress in restoring speech by assessing signals from motor structures^98–100^, and integrating planning signals from the areas we identified may lead to further improvements in decoding performance. Furthermore, our rigorous classification of language-related cognitive and sensorimotor brain dynamics informs a range of communication disorders thought to result from disordered planning^5,92,101,102^.

## ACKNOWLEDGEMENTS

We thank members of the Long laboratory as well as Margot Elmaleh, Jelena Krivokapić, and Jeremy Yeaton for helpful feedback on early versions of this manuscript. We also thank Haiming Chen, Christopher Garcia, Matthew Howard III, Kenji Ibayashi, Kirill Nourski, Ariane Rhone, Andrea Rohl, and Beau Snoad for help with data collection.

This research was supported by R01DC019354 (M.A.L.), R01DC015260 (J.D.W.G.), the Simons Collaboration on the Global Brain (M.A.L.), and a Career Awards at the Scientific Interface fellowship from the Burroughs Wellcome Fund (G.A.C.).

## AUTHOR CONTRIBUTIONS

Conceptualization: GAC, MM, MAL; Data curation: GAC, CKK, FT; Formal analysis: GAC, MM; Funding acquisition: GAC, JDWG, MAL; Investigation: GAC, CKK, JDWG; Methodology: GAC, MM, CKK, FT, JDWG; Project administration: GAC, JDWG, MAL; Resources: GAC, JDWG, MAL; Software: GAC, CKK, FT; Supervision: GAC, JDWG, MAL; Validation: GAC, CKK, JDWG; Visualization: GAC, MAL; Writing – original draft: GAC; Writing – review & editing: GAC, MM, CKK, FT, JDWG, MAL

## METHODS

### Resource Availability

Requests for additional information and resources should be directed to the lead author, Michael A. Long.

### Study participants

Twenty-two patient-volunteers undergoing surgical treatment at the University of Iowa Hospitals and Clinics for medically intractable epilepsy, brain tumors, or movement disorders were participants in this study. During their treatment, neural activity was recorded using electrocorticography (ECoG) electrodes and/or intracerebral stereoelectroencephalography (SEEG) depth electrodes either chronically (i.e., for seizure focus determination) or acutely (i.e., during awake craniotomy for tumor removal/epilepsy treatment or during a deep-brain stimulation electrode implant procedure). No participants were excluded based on language lateralization, but the majority were likely left lateralized for language function based on Wada testing, right-handedness, and/or the occurrence of left hemisphere speech arrest sites during intraoperative language mapping (n = 19). Finally, all participants were native speakers of English. Additional demographic details and clinical information is available in Table S1. All study participants consented to research, and the University of Iowa Institutional Review Board approved all procedures.

### Data acquisition and processing

#### Acquisition

For awake craniotomy patients, local field potentials were recorded using subdural ECoG arrays manufactured by Ad-Tech Medical or PMT Corporation. Signals were amplified and sampled at 2034.5 Hz using a multichannel amplifier and digital acquisition system (PZ2 or PZ5 preamplifier with an RZ2 processer; Tucker-Davis Technologies). For chronically implanted epilepsy patients, electrical signals from subdural ECoG arrays or SEEG intracerebral depth electrodes (Ad-Tech) were recorded at 2000 Hz with a multichannel amplifier and digital acquisition system (Atlas system, Neuralynx). In both contexts, analog input channels synchronized with the neural recordings additionally acquired the output of 1-3 microphone(s) which captured the speech acoustics of the experimenter and participant. Input channels were typically sampled at 48,828 Hz by the TDT system and 16,000 Hz by the Neuralynx system and downsampled offline. In addition to the electrical signals, a video of the participant was often acquired at 24 fps for 20 of 22 experiments. The video was synced to the electrophysiological data after the experiment and provided an additional high-quality audio recording channel sampled at 48 kHz.

#### Electrophysiological signal processing

Raw ECoG and SEEG signals were first visually inspected and any dead channels or channels displaying artifact were rejected. Data were then preprocessed by removing stationary and nonstationary line noise using adaptive thresholding applied to coefficients of the demodulated band transform^51^ with a bandwidth parameter of 0.25 Hz. The task period of recordings (five seconds before the onset of the first CI trial until 5 seconds after the offset of the last CI trial) was then isolated and any electrodes not on the lateral, ventral, or superior temporal plane surface were excluded from further analysis. Data were then prefiltered at 3 Hz (FIR high-pass filter) and each grid was common average re-referenced. The high gamma (HG) amplitude for each electrode was next calculated by bandpass filtering the signal at 70-150 Hz (FIR bandpass), performing a Hilbert transform, extracting its absolute value, and smoothing with quadratic regression in 500 ms moving windows. The resulting HG signals were then resampled to 500 Hz and any long timescale drift was removed by subtracting the moving average of each channel calculated across 5 minutes. Finally, all HG signals were z-scored across the entire task period.

### Anatomical reconstructions

#### Electrode localization

Depending on clinical constraints, subdural electrode locations were localized using a combination of peri-operative magnetic resonance, peri-operative computed tomography, intraoperative fluoroscopy, and/or high-resolution photographs of the craniotomy. All localization procedures are explained in detail in our previous publication^36^.

#### Electrode coregistration

All electrode locations were first determined on cortical surface renderings in RAS coordinate space. Preoperative T1 images were then non-linearly coregistered to an MNI-aligned template brain (CIT168 template)^103^ using symmetric diffeomorphic registration in the ANTs toolbox^104^. Finally, the electrode locations were rendered on canonical cortical surfaces (e.g., Figure 1A) by plotting their coregistered coordinates on the gyral surface of the MNI152 brain.

#### Behavioral task

Participants performed the Critical Information (CI) task, which entailed listening to a set of simple questions read by an experimenter and responding as quickly as possible. The CI questions were adapted from an established Dutch stimulus set^105,106^ and used in our previous work^35,36^. Each CI question contains a single word (i.e., the critical information) which is necessary for answering the question (e.g., Figure 1B) and requires participants to generate the antonym of common words (e.g., The opposite of soft is what frequent word?), name the animal corresponding to vocalization (What animal, who oinks, is commonly seen on farms?), and/or report the number of body parts typically found on humans (e.g., How many arms does a person have?). Questions were presented in randomized order and, in some experiments, were interleaved with trials from other tasks that are not analyzed in this study.

To determine the timing of all questions, critical information, and answers, the audio acquired with the electrophysiological acquisition system and/or video camera was annotated and timestamped by a trained phonetician (G.A.C.). Only trials where the participant answered correctly and without a hesitation (e.g., ‘uh’, ‘um’, or other disfluencies) were analyzed.

#### Quantification and Statistical Analyses

Details regarding electrophysiological signal analysis and statistical testing can be found below. Statistical details can be found in figure legends and the Results section (e.g., statistical tests used, exact value of n, what n represents). Summary statistics are reported as mean ± one standard deviation unless otherwise noted. Normal distribution of data was not assumed.

### Detection of significant neural responses

We used a resampling approach to detect electrodes that were significantly modulated during the CI task (Figure S1). Because CI task performance can vary widely within and across participants, our first step was to restrict our analysis to trials with comparable behavior by instituting a maximum reaction time cutoff. However, responses are initiated systematically faster when the CI is presented early in the question (early trials; Figure 1B, right) rather than at the end^32,36,105^ (late trials; Figure 1B, left), thus we applied separate maximum reaction time limits for these trial types. Specifically, early trials with reaction times greater than 1 second and late trials with reaction times greater than 1.275 seconds were excluded from analysis, which approximated the 75^th^ percentile of both distributions (1.001 seconds and 1.232 seconds, respectively) (Figure S1A). Following the above behavioral standardization step, we next calculated the average HG responses for all electrodes that were not located over seizure foci or a tumor when aligned to four task timepoints (Figure 1B) ± 2 seconds after rejecting any trials containing artifact (i.e., a period surpassing 9z in amplitude) (Figure S1B). These timepoints were selected to enable analysis windows that sampled most of the task while not overlapping.

To determine periods of significant activation and suppression in the average electrode responses, we then calculated the shuffled average response of each electrode aligned to random timepoints in the task period of the recordings while preserving trial numbers. This process was repeated over 1000 iterations to generate a null distribution of average response amplitude and calculate its 95% confidence interval, which allowed us to identify periods of the actual response that fell outside of the null distribution (Figure S1C). We next assessed whether these putative active and suppressed periods were significantly longer in duration than chance. To do so, we assessed the distribution of analogous periods in the shuffled responses, thus generating null distributions for the durations of putative active and suppressed periods (Figure S1D). Finally, any putative periods of activation or suppression whose duration was higher than the 95^th^ percentile value of the corresponding null distribution were considered significantly active or suppressed periods, respectively (Figure S1E).

Finally, any electrode that displayed a period of significant activation or suppression within specific epochs around at least one of the four alignment points was deemed to be significantly modulated during the CI task. These epochs were designed to control for differences between CI trial types and CI question presentation across participants while minimizing overlap between analysis windows. The exact timing of these epochs was based on the structure and timing of the analyzed trials, and were designated as follows: 1) 0.25 seconds before to 1 second after question onset in early trials, 2) 0.25 seconds before to 1.5 seconds after CI onset in early trials, and 3) 1.65 seconds before to 1.25 seconds after answer onset in both early and late trials.

### Clustering of neural responses

To test whether electrode responses formed functional clusters, we first performed dimensionality reduction on the signals from all significantly modulated electrodes using principal component analysis (PCA). To do so, we specifically concatenated the signals from each modulated electrode in the four task epochs described above after resampling such that each epoch contributed an equal number of samples; these response vectors were then z-scored and input into the ‘pca’ function in MATLAB with default settings. We then used four methods (k-means, k-medoids, Gaussian mixture modelling, and Ward’s hierarchical clustering) to cluster the electrodes according to their coefficient for the first 3 PCs. For each method, we assessed clustering performance by computing the empirical silhouette score and the Calinski-Harabasz (C.-H.) criterion using MATLAB’s ‘evalclusters’ function while assuming 2-10 clusters and repeating this process over 500 iterations for all methods except Ward’s hierarchical clustering (which is deterministic). Finally, because the k-means algorithm when assuming 4 clusters resulted in the highest median silhouette scores and C.-H. criteria, we used the k-means optimal clustering solution over the 500 iterations to define the electrode clusters in subsequent analyses.

We further assessed the clustered structure of the modulated electrode responses by performing a Parallel Analysis-based method^107^ with the same PCA and clustering analysis on shuffled responses (Figure S2), which also allowed us to empirically define the number of PCs to use for our final clustering analyses (i.e., 3 PCs; Figure S2A). For this analysis, we randomly reassigned the responses in each window such that each electrode possessed a random combination of real responses aligned to the four timepoints. This randomization approach was repeated 1,000 times, and we performed k-means clustering over 500 iterations while recording the empirical silhouette score and C.-H. criterion when assuming 2-10 clusters. For each shuffled data set, the optimal number of clusters was defined according to median silhouette score and the optimal clustering solution was defined as the iteration that resulted in the highest overall silhouette score when assuming the optimal number of clusters.

### Behavioral Analysis

To analyze how electrodes were modulated during different behavioral phases of the CI task, we first calculated the median duration of four epochs in early and late trials separately: question onset to CI onset, CI onset to question offset, question offset to answer onset, and answer onset to answer offset. This allowed us to determine the time course of ‘typical’ trials that were normalized to these median values across participants (e.g., Figure S3A) and thus adopt analysis time windows corresponding to particular task phases. We then assessed the percentage of electrodes whose average responses exhibited significantly elevated and suppressed activity for at least 50 ms in the following analysis windows (see numbered bars in Figure S3A): 1) 0 to 1.5 or 0 to 0.5 seconds after question onset in late and early trials, respectively, 2) 0 to 0.5 or 0 to 1.7 seconds after CI onset in late and early trials, respectively, 3) 0.1 to 0.5 or 0.1 to 0.3 seconds after question offset in late and early trials, respectively, and 4) 0 to 0.6 seconds after answer onset in early or late trials (Figure S3B). Finally, we also calculated each electrode’s peak activity (i.e., 90^th^ percentile value of average response) and suppression (i.e., 10^th^ percentile value of average response) in each of these four windows (Figure S3C).

The same task duration normalization procedure was performed using behavioral data from participant 494L to extract the data presented in Figure 4C (see Figure 4B for ‘typical’ early trial in this participant). For each electrode in the example grid, we calculate the mean response amplitude in the following early trial windows: 1) -0.35 to -0.5 seconds aligned to CI onset, 2) 0.05 to 0.35 seconds aligned to CI onset, 3) 0.5 to 0.8 seconds aligned to CI onset, 4) -0.35 to -.005 aligned to answer onset, 5) 0.1 to 0.5 seconds aligned to answer onset, and 6) 0.6 to 0.9 seconds aligned to answer onset.

### Anatomical analysis

To test whether response clusters were spatially structured on the cortical surface, we examined if nearby electrodes came from the same cluster. For each modulated electrode, we first calculated the proportion of modulated electrodes with matching cluster assignments (‘same-cluster’ electrodes) that were located within a 1-cm radius on individual cortical surfaces and when the data was pooled across participants on the coregistered cortical surface. We then repeated the same procedure but randomized cluster identity across all electrodes (i.e., including unmodulated electrodes) within individual participants or across the entire data set over 1,000 iterations, which generated null distributions for each modulated electrode. Finally, we assessed significance by determining whether the actual median proportion of nearby same-cluster electrodes in the within-participant and pooled data sets fell outside the range of median values observed in the corresponding null distribution.

We then identified areas of the cortical surface which contained significantly more electrodes of each cluster than would be expected by chance using a spatial permutation test. We did so by dividing the MNI152 brain into 1-cm cubic voxels with 90% overlap (centers of each voxel were 1 mm apart in the x, y, and z dimensions) and counting the number of electrodes from a given cluster that were located within all voxels. We then generated the null distribution of electrode counts by shuffling cluster labels across all electrode coordinates (i.e., so that cluster labels were randomly assigned to electrodes) over 1,000 iterations and counting the number of electrodes from the cluster that were located within all voxels. To assess significance, we then identified all voxels whose actual number of electrodes exceeded the 95^th^ percentile of the null distribution; to avoid spurious results, any voxels containing two or fewer electrodes from a cluster were not subjected to significance testing. To visualize the location of voxels containing significantly more electrodes of a given cluster than expected by chance (Figure 3E-H), we plotted the location of all electrodes from that cluster on the MNI152 canonical cortical surface.

### Nonnegative matrix factorization

We performed nonnegative matrix factorization (NMF) on the responses of all planning and motor class electrodes aligned to critical information and answer onset in early CI trials, respectively. To do so, we first selected the number of factors to assume for each response class by assessing root mean square reconstruction error (RMSE) and variance accounted for (VAF). We then performed NMF (1,000 maximum iterations and one replicate, using MATLAB’s built-in multiplicative update algorithm) on all responses of each class separately in the same time windows used for significant response detection. This process was repeated 1,000 times when assuming 1-10 factors, which allowed us to determine the distribution of VAF (Figure S4A) and RMSE values (Figure S4B). The distributions of these metrics across different numbers of factors revealed a linear decay phase where assuming more factors did not lead to meaningful increases in VAF or RMSE. To estimate the onset of this linear decay phase, we examined the change in slope between points in the RMSE plot by performing sequential linear fits between the values when assuming one factor and all other factors (Figure S4B). We found that the change in slope when assuming an increasing number of factors peaked (Figure S4C) at the onset of the linear decay phase in the RMSE data and corresponded to an ‘elbow’ point in the distribution (Figure S4B). We therefore selected the number of factors corresponding to this peak for further analysis, which corresponded to the models assuming the fewest factors that accounted for a minimum of 90% VAF (Figure S4A). We then performed a final NMF analysis for each response class assuming the selected number of factors (100,000 maximum iterations and 100,000 replicates, using MATLAB’s built-in alternating least-squares algorithm) and used the optimal model observed across our first 1,000 replicates as an initialization point.

To assess whether electrodes were clustered according to their NMF factor weights, we clustered planning and motor class electrodes separately using k-means clustering over 1,000 iterations while assuming 1-10 clusters. We then plotted the distribution of empirical silhouette scores (Figure S4D) and C.-H. criteria (Figure S4E) for each number of clusters, and we did not observe strong clustering for either response class (e.g., Figures 1 and S2).

Finally, we investigated whether electrodes were anatomically organized using a modified version of the spatial permutation test described above. Rather than counting the number of electrodes within 1-cm^3^ voxels, we calculated the median NMF factor weight in all voxels containing at least 3 planning or motor class electrodes; this process was performed separately for each factor. To generate a null distribution of this metric, we shuffled factor weights across electrodes of the relevant class and calculated the median weight over 1,000 iterations. Any voxel whose observed median factor weight surpassed the 95^th^ percentile of shuffled median values was defined as significantly overweighted for that factor. To visualize the location of significantly overweighted voxels, the location of electrodes within such voxels which possessed factor weights higher than the median across electrodes was plotted on the MNI152 canonical cortical surface. On the MNI152 brain, we also plotted the density of these electrodes relative to all electrodes analyzed in a 7.5 mm radius (smoothed with a Gaussian kernel). Specifically, proportional density at each vertex of the MNI152 surface mesh was determined by summing over the kernel values with respect to included electrode location and dividing the result by the same summation over all electrode locations. Values were plotted only at vertices with 3 or more electrodes within 7.5 mm. The same procedure was used to visualize the density of electrodes from each response class (Figure S3D-G).

### Statistical testing

All analyses, including permutation tests, were performed in MATLAB (Mathworks, Natick, MA) using a combination of built-in functions and custom scripts.

