## Supplemental Figures and Table for "Neural activity flows through cortical subnetworks during speech production"

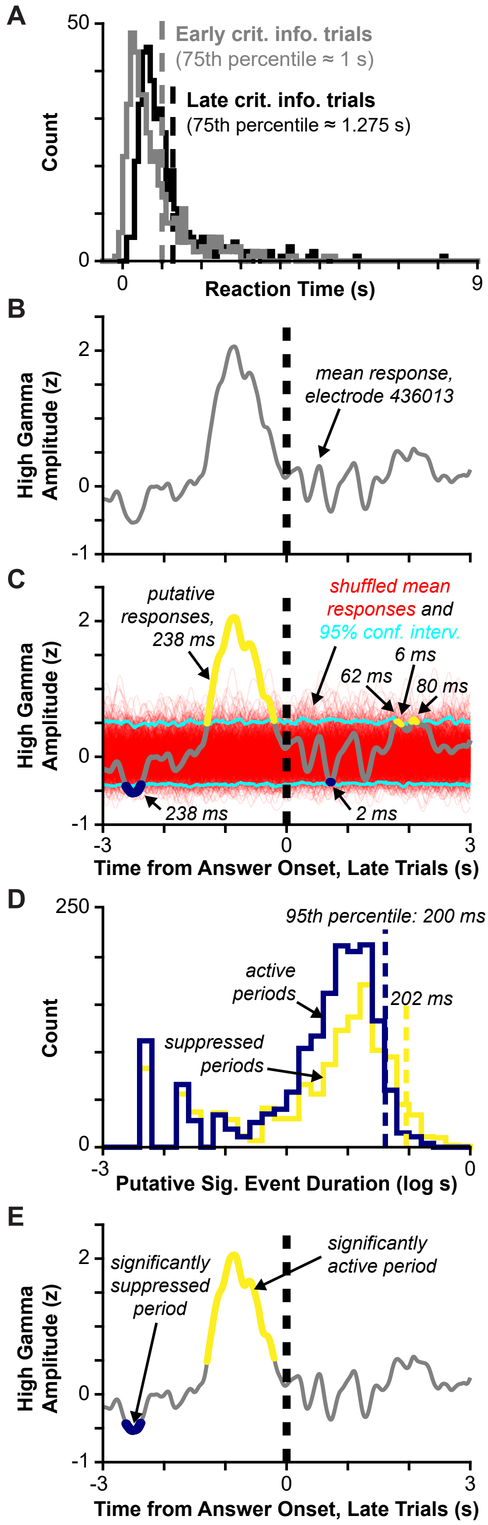


***1) Behavioral thresholding*** to control for differences in task performance and trial structure across participants.

**2)** Calculate ***average electrode response*** across trials aligned to behaviorally relevant timepoints.

**3)** Shuffle alignment points to determine ***amplitude null distribution***.

**4)** Determine ***duration null distribution*** using the timing of all putative significant events in shuffled responses.

**5)** Identify **periods of significant activation and suppression** in actual average response.

**Figure S1. Method for responsive electrode detection.**
(A) Reaction time distribution for all early and late CI trials pooled across participants with maximum reaction time limits indicated.
(B) Average response (i.e., high gamma amplitude) of an example electrode aligned to participant answer onset in late CI trials.
(C) Same as (B), but with 1,000 shuffled responses (i.e., average responses aligned to randomized timepoints) and the 95% confidence interval of the shuffled response amplitude overlaid. Any periods of the actual mean response above (in yellow) or below (in dark blue) the confidence interval are considered 'putative responses' and are labeled with their durations.
(D) For the presented electrode, the distribution of putative active and suppressed durations observed in shuffled responses (i.e., periods of the red responses in (C) that are above and below the 95% confidence interval, respectively) with the 95$^{th}$ percentile values indicated.
(E) For the example electrode, putative active (yellow) and suppressed (dark blue) periods in the average response whose duration surpasses the 95$^{th}$ percentile values indicated in (D); this electrode is significantly suppressed or active in these periods.

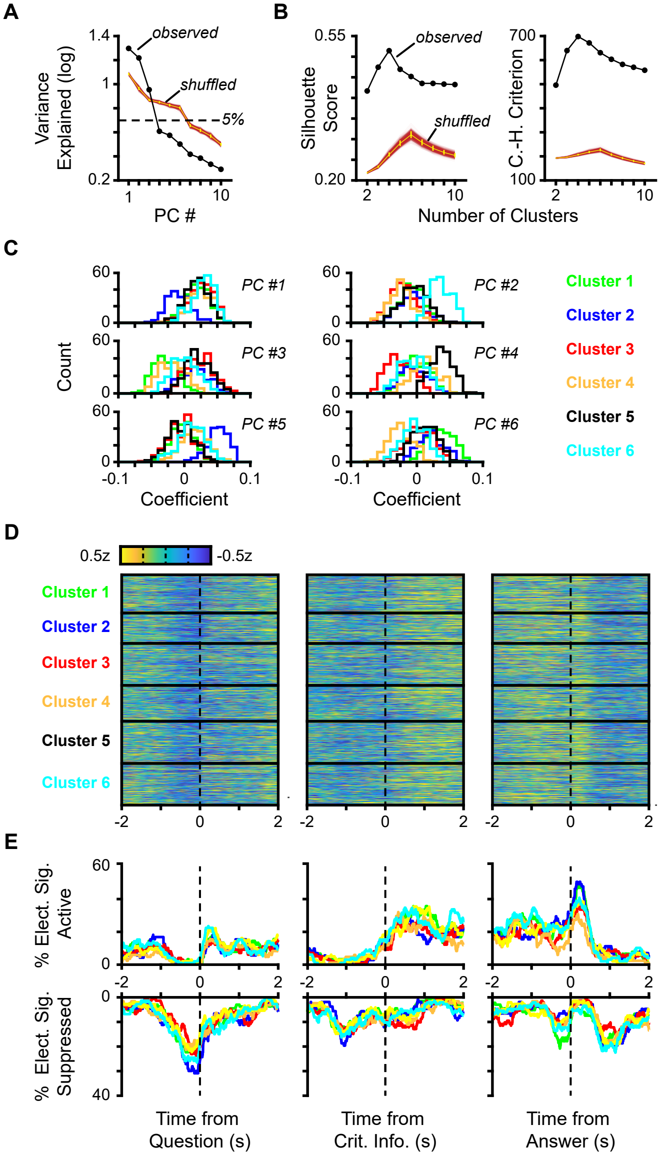

A
B
C
D
E
1.4
observed
1
shuffled
5%
0.2
Variance Explained (log)
PC #
0.55
Silhouette Score
0.20
700
C.-H. Criterion
100
Number of Clusters
PC #1
PC #2
PC #3
PC #4
PC #5
PC #6
Count
Coefficient
Cluster 1
Cluster 2
Cluster 3
Cluster 4
Cluster 5
Cluster 6
0.5z
-0.5z
% Elect. Sig. Active
% Elect. Sig. Suppressed
Time from Question (s)
Time from Crit. Info. (s)
Time from Answer (s)

**Figure S2. Additional clustering analyses.**
(A) Parallel Analysis: Amount of variance explained by each principal component (PC) in the observed data set and when responses within each analysis window are randomly shuffled across significantly modulated electrodes (see Methods); for the shuffled data, thin lines indicate the values for each iteration, the yellow line indicates the median across iterations, and yellow error bars indicate the $10^{th}$ and $90^{th}$ percentile values.
(B) Median silhouette scores (left) and Calinski-Harabasz (C.-H.) criteria (left) for k-means clustering for the observed data set and across 1,000 trials of response shuffling; color scheme is the same as in (A).
(C) For the optimal clustering solution across all iterations of response shuffling, the histograms depicting the distribution of PC coefficients for each response cluster.
(D) For the optimal clustering solution across all iterations of response shuffling, the average electrode response for each activity cluster aligned to question onset in late trials (left), critical information onset in early trials (middle), and answer onset in early trials (right).
(E) For the same alignment points, the percentage of electrodes within each cluster displaying either significantly elevated (top) or suppressed (bottom) activity over time.

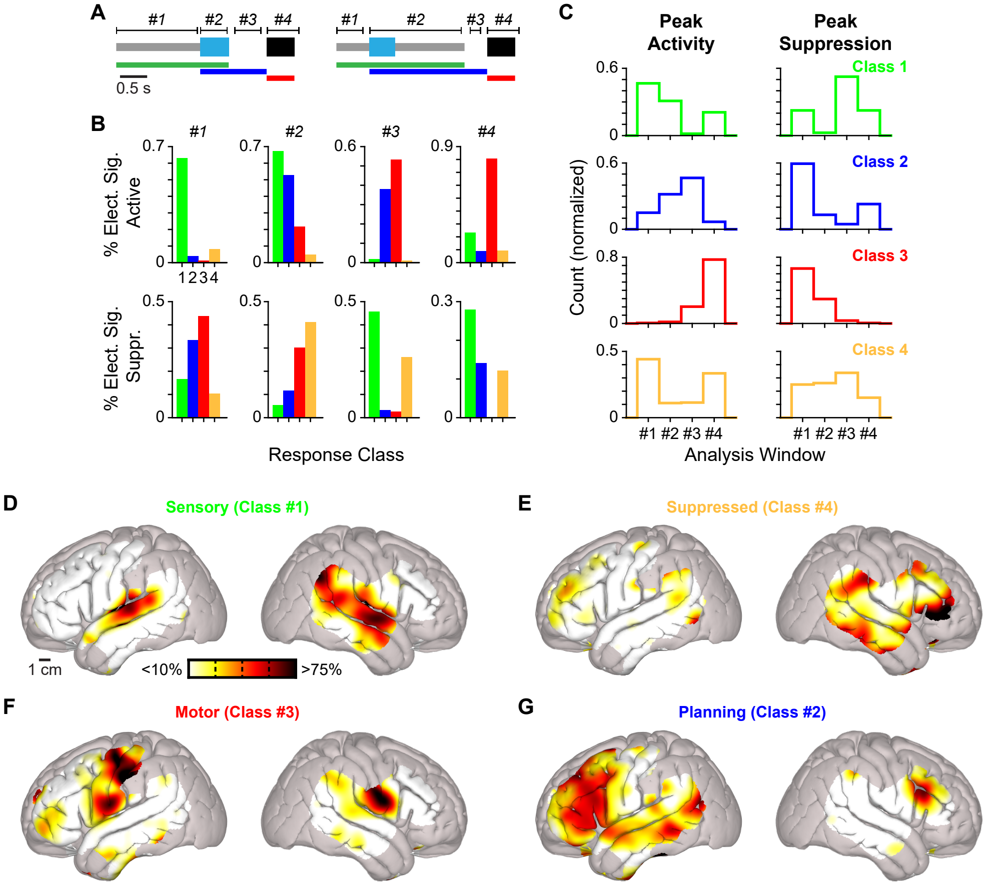

A
#1 #2 #3 #4
#1 #2 #3 #4
0.5 s
B
#1 #2 #3 #4
% Elect. Sig. Active
% Elect. Sig. Suppr.
1234
Response Class
C
Peak Activity
Peak Suppression
Class 1
Class 2
Class 3
Class 4
Count (normalized)
#1 #2 #3 #4
Analysis Window
D
Sensory (Class #1)
1 cm
<10%
>75%
E
Suppressed (Class #4)
F
Motor (Class #3)
G
Planning (Class #2)

**Figure S3. Additional functional and anatomical analyses.**
(A) Diagrams of a late (left) and early (right) trial with sensory, planning, and motor phases indicated using colored bars and analysis windows numbered; color scheme is identical to Figure 1B and trial duration is normalized (see Methods).
(B) For each response class, the percentage of electrodes that were significantly active (top) and suppressed (bottom) for at least 50 ms within the task analysis windows denoted in (A) for late and/or early trials; activity in window #1 is aligned to question onset, in window #2 is aligned to CI onset, in window #3 is aligned to question offset, and in window #4 is aligned to answer onset.
(C) For every response class, the percentage of electrodes displaying peak activity or suppression levels (see Methods) in each analysis window.
(D-G) Heatmaps illustrating the density of all electrodes in each response classes (spatially smoothed with 7.5 mm Gaussian kernel; see Methods).

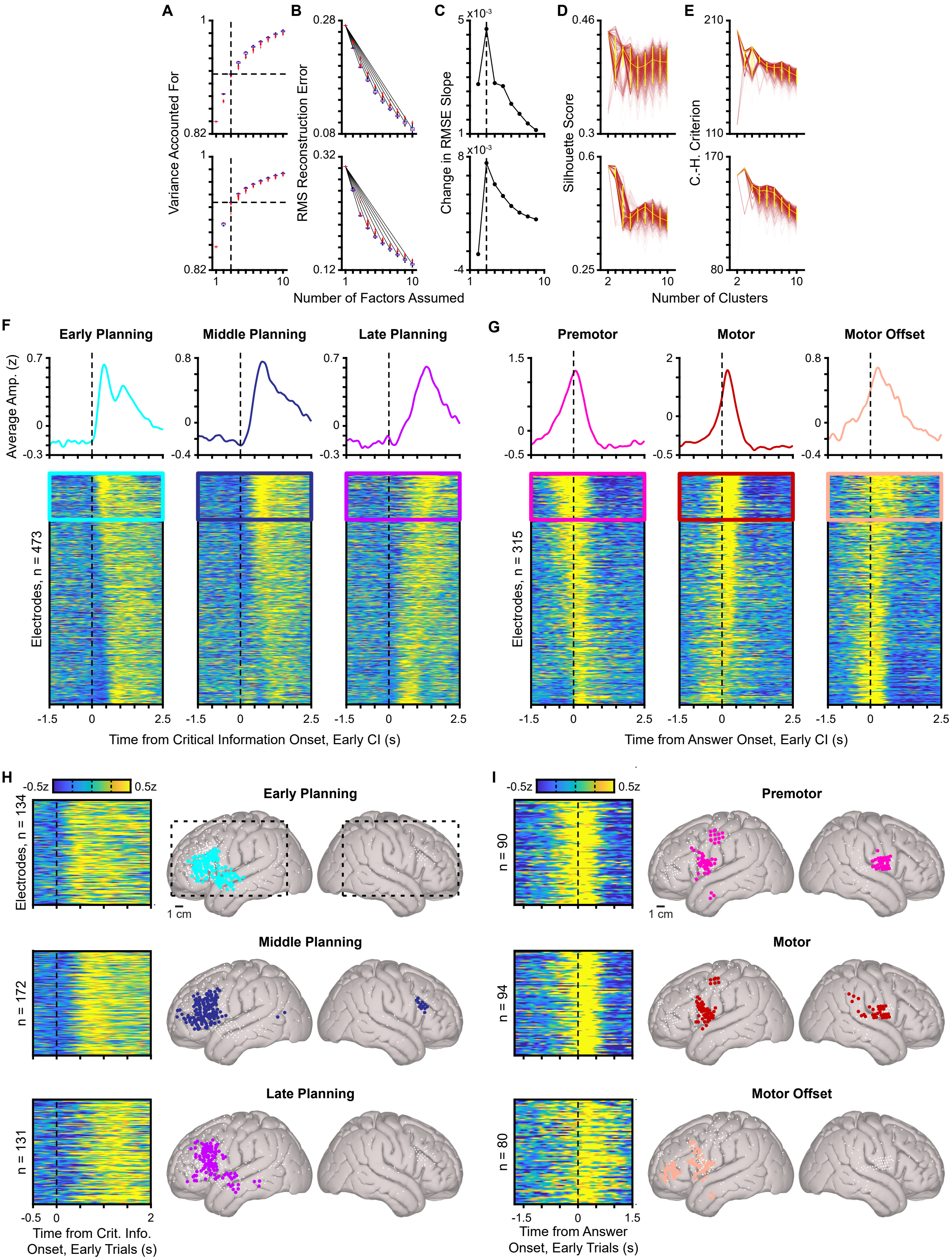

A
B
C
D
E
Variance Accounted For
RMS Reconstruction Error
Change in RMSE Slope
Silhouette Score
C.-H. Criterion
Number of Factors Assumed
Number of Clusters
F
G
Early Planning
Middle Planning
Late Planning
Premotor
Motor
Motor Offset
Average Amp. (z)
Electrodes, n = 473
Electrodes, n = 315
Time from Critical Information Onset, Early CI (s)
Time from Answer Onset, Early CI (s)
H
I
-0.5z
0.5z
Electrodes, n = 134
n = 172
n = 131
n = 90
n = 94
n = 80
1 cm
Time from Crit. Info. Onset, Early Trials (s)
Time from Answer Onset, Early Trials (s)

**Figure S4. Additional NMF-related analyses.**
(A) Boxplots depicting the variance accounted for (VAF) when assuming a different number of factors across 1,000 iterations of nonnegative matrix factorization (NMF) for planning (top) and motor (bottom) class responses; dashed lines indicate the number of factors selected for the analyses in Figure 4.
(B) Same as (A), except for root mean square error (RMSE); solid black lines indicate linear fits between the RMSE error distribution when assuming one factor and all other numbers of factors.
(C) Difference between the slope values for the linear fits in (B), with data for planning responses at top and motor responses at bottom; dashed lines indicate the maximal change in slopes and the number of factors selected for the analyses in Figure 4.
(D) Silhouette scores when planning (top) and motor (bottom) class responses are clustered according to their NMF weights (i.e., Figure 4B,C, left) over 1,000 iterations of k-means clustering when assuming 1-10 clusters; thin lines indicate the values for each iteration, the yellow line indicates the median across iterations, and yellow error bars indicate the $10^{th}$ and $90^{th}$ percentile values.
(E) Same as (D), except for Calinski-Harabasz (C.-H.) criteria.
(F) Average responses for all planning class electrodes ordered by descending weights for all 3 NMF factors; at top, the mean response across the most highly weighted 20% of electrodes (indicated by box).
(G) Same as F, but for motor class electrodes.
(H-I) The location of all planning (H) and motor (I) electrodes within voxels that are weighted significantly higher for each component response profile than expected by chance ($p < 0.05$, permutation test) on canonical cortical surfaces (right); electrodes with a weight higher than the median value are indicated with large colored points and their responses are shown at left, all other electrodes within a given response class are indicated with white points. Dashed boxes indicate the cortical surfaces regions depicted in Figure 4F.

**Table S1. Participant demographic and clinical information.**

| Number | Age | Sex | Recording Type | Coverage | Language Dominance (Method) | Handedness | Pathology/Diagnosis | Tumor Location/Seizure Loci |
|---|---|---|---|---|---|---|---|---|
| 436 | 58 | F | Intraoperative | Left lateral | Unknown; speech arrest induced by stimulation of left precentral gyrus | right | Grade II oligodendroglioma (tumor) | Left middle frontal gyrus |
| 442 | 35 | F | Chronic | Left lateral, ventral, superior temporal plane | LH (Wada testing) | right (+70) | Epilepsy | Multifocal onset |
| 456 | 30 | M | Chronic | Right lateral, ventral, superior temporal plane | LH (Wada testing) | left (-40) | Epilepsy | right middle hippocampus and amygdala; inferior area of the supramarginal gyrus at the proximity of the angular gyrus |
| 460 | 51 | M | Chronic | Left lateral, ventral, superior temporal plane | LH (Wada testing) | right | Epilepsy (gliosis) | Left mesial temporal lobe |
| 463 | 65 | M | Intraoperative | Left lateral | Unknown; speech arrest by stimulation at the level of the left supramarginal gyrus. | ambidextrous | Glioblastoma | Left parahippocampal gyrus |
| 472 | 32 | M | Intraoperative | Left lateral | Unknown | right | Vascular malformation, focal cortical dysplasia TIIIc | Left temporal pole |
| 477 | 24 | F | Chronic | Left lateral | LH (Wada testing) | right (+80) | Epilepsy (mild gliosis) | Left parietal |
| 486 | 53 | M | Intraoperative | Left lateral | LH (Wada testing) | right | Glioblastoma | Left inferior frontal gyrus |
| 494 | 30 | M | Intraoperative | Left lateral | LH (Wada testing) | right (+70) | Hippocampal sclerosis (epilepsy) | Left hippocampus |
| 510 | 54 | F | Intraoperative | Left lateral | Unknown; motor speech arrest area in the pars opercularis of the left inferior frontal gyri following stimulation. | right | Anaplastic Oligodendroglioma | Left inferior frontal gyrus |
| 516 | 58 | F | Intraoperative | Left lateral | Unknown; speech arrest induced by stimulation of anterior to the left motor cortex (potentially in inferior frontal gyrus) | right | Meningioma, WHO grade I | Left partietal lesion (posterior to the post central gyrus) |
| 528 | 22 | F | Intraoperative | Right lateral | LH (Wada testing) | right | Epilepsy | Right mesial temporal lobe |
| 532 | 42 | F | Chronic | Right lateral, ventral, superior temporal plane | LH (Wada testing) | right(+100) | Epilepsy | right temporal or frontal temporal area |
| 585 | 38 | F | Chronic | Left lateral, ventral, superior temporal plane | LH (Wada testing) | right | Epilepsy | left middle hippocampus |
| 595 | 37 | F | Intraoperative | Left lateral | bilateral language function (Wada) | left | Epilepsy | left anterior temporal lobe |
| 671 | 71 | M | Intraoperative | Left lateral | Unknown | right | Parkinson's Disease | N/A |
| 682 | 68 | M | Intraoperative | Left lateral | Unknown | right | Parkinson's Disease | N/A |
| 706 | 56 | F | Intraoperative | Left lateral | Unknown | right | Essential Tremor | N/A |
| 716 | 51 | M | Intraoperative | Left lateral | Unknown; dysarthria induced by stimulation of left precentral gyrus | left | Essential Tremor | N/A |
| 750 | 61 | M | Intraoperative | Left lateral | Unknown | right | Parkinson's Disease | N/A |
| 758 | 65 | M | Intraoperative | Left lateral | Unknown | left | Essential Tremor | N/A |
| 768 | 64 | F | Intraoperative | Left lateral | Unknown | right | Essential Tremor | N/A |
